# Genetic conversion of a split-drive into a full-drive element

**DOI:** 10.1101/2021.12.05.471291

**Authors:** Gerard Terradas, Jared B. Bennett, Zhiqian Li, John M. Marshall, Ethan Bier

## Abstract

Gene-drive systems offer an important new avenue for spreading beneficial traits into wild populations. Their core components, Cas9 and guide RNA (gRNA), can either be linked within a single cassette (full gene drive, fGD) or provided in two separate elements (split gene drive, sGD) wherein the gRNA-bearing element drives in the presence of an independent static source of Cas9. We previously designed a system engineered to turn split into full gene drives. Here, we provide experimental proof-of-principle for such a convertible system inserted at the *spo11* locus, which is recoded to restore gene function. In multigenerational cage studies, the reconstituted *spo11* fGD cassette initially drives with slower kinetics than the unlinked sGD element (using the same Mendelian *vasa-Cas9* source), but eventually reaches a similar level of final introgression. Different kinetic behaviors may result from transient fitness costs associated with individuals co-inheriting Cas9 and gRNA transgenes during the drive process.

## Introduction

CRISPR-based gene drives offer novel approaches for vector control by transforming genetic structures of wild insect populations ^1–5^. Linked gene drives (or full gene drives, fGD) carry gene cassettes that encode the bacterial Cas9 endonuclease and a guide RNA (gRNA) sequence that directs the Cas9/gRNA ribonucleoprotein complex to cleave the genome at the site of cassette insertion ^3–5^. Upon cleavage of the target site on a homologous chromosome, the gene-drive cassette is copied into the double stranded break (DSB) by homology-directed repair (HDR) using the drive-bearing homologous chromosome as a repair template. If this directional gene conversion process is efficient, the drive allele will be inherited in super-Mendelian fashion (>50%). Genes encoding beneficial factors can be linked to fGD cassettes to spread these traits into susceptible populations within a few generations ^3^.

Gene-drive technologies can be used either to modify or suppress insect disease vector populations depending on their design. Modification drives ^3,6,7^ carry a beneficial cargo to reduce the vector capacity of the insect, while suppression drives ^5,8,9^ endeavor to bias the inheritance of deleterious traits that will ultimately either kill or sterilize the insect. In addition to autonomously acting fGDs in which Cas9 and gRNA are encoded within a unitary cassette, so-called split gene-drives (sGD) have also been developed ^10–12^ in which the copying cassette carries only the gRNA component, with Cas9 supplied from a second genomic site. In such bi-partite arrangements, the gRNA-carrying cassette is copied in the presence of the Cas9 source while the Cas9-encoding element is static (i.e., transmitted in a standard Mendelian fashion). In the absence of Cas9, however, the gRNA-bearing split-drive element also is inherited at Mendelian frequencies.

Because the copying component of a split gene drive spreads in an additive rather than exponential fashion ^4^, these elements have been deemed safer and offer the means for more localized alterations of target populations ^13^. The split configuration also provides greater flexibility for experimental analysis of the various CRISPR components including different promoters driving Cas9 expression and comparison of alternative gRNAs ^12,14^. These advantages obviate the need for developing new full gene drives to test each pair of Cas9 and gRNA, which is time-consuming, expensive, and requires a higher level of biological confinement. A modification of the traditional split gene drive design is the trans-complementing gene drive (tGD) ^11^, in which the gRNA cassette carries a second gRNA that targets cleavage of the genome at the site where the Cas9 element is inserted. When kept as separate stocks, the Cas9 and gRNA encoding elements are inherited in a Mendelian fashion, but when they are combined (e.g., by crossing stocks carrying the elements to each other) they behave as a conjoined full gene drive system. Although there are many advantages associated with the tGD system, it also is difficult to confine once assembled as a dual unit. Furthermore, the requirement of driving two separate elements magnifies the effect of drive-resistant mutations generated by the alternative non-homologous end-joining (NHEJ) pathway ^15^ that can arise at either gRNA cleavage site, potentially limiting the drive kinetics and final fraction of the population carrying both elements.

Our group recently analyzed a set of sGDs inserted into essential genes carrying functional, recoded cDNA sequences that restore functionality of the endogenous loci upon allelic cleavage and conversion ^12^. Such recoded sGDs minimize the production of non-functional NHEJ alleles that are resistant to Cas9 cleavage by dominantly eliminating such alleles transmitted by females through a process referred to as sterile/lethal mosaicism ^6,16^. This latter process is based on maternal inheritance of Cas9/gRNA complexes that act on the paternal allele to mutate it in a sufficient fraction of cells in progeny to either kill (lethal mosaicism) or sterilize (sterile mosaicism) such individuals. Although unexploited in the previous study ^12^, the sGD constructs included another design feature that permits facile genetic transfiguration of the sGD into an fGD. This conversion system relies on a set of Cas9 sources that carry sGD homology arms flanking the Cas9 cassette and a gRNA (gRNAHack) that can cleave a synthetic target site within the sGD element. When these Cas9 sources (inserted into AttP sites in different genomic locations) are crossed to the sGD element, the gRNAHack mediates cleavage of the sGD resulting in the insertion of the Cas9 cargo into the recipient element. Here, we provide proof-of-concept for such conversion of a recoded split drive into a full drive inserted into the *spo11* locus, which is required for fertility in both sexes. This strategy for converting a sGD into a fGD follows the logic of the homology-assisted CRISPR knock-in system (or HACK) ^17^, efficiently used to generate DNA double-stranded breaks and replace GAL4 cassettes in *Drosophila*. This flexible genetic swap system can rapidly convert an optimally performing sGD into an fGD and permits direct comparison of the performance of split versus full-drive systems that are inserted at identical genomic sites and are powered by the same promoter-Cas9 transgene. We evaluate the performance of a converted (or hacked) fGD in both single generation crosses and cage competition experiments and compare its performance to that of the sGD described in previous experiments ^12^. We find that while the fGD displays slower initial drive kinetics, it catches up to the sGD and achieves a similar level of final introgression into the population. The delayed drive of the fGD element may result from a transient fitness cost we have previously described in sGD experiments that is associated with co-inheritance of Cas9 and gRNA transgenes in presence of cleavage-sensitive target alleles ^12^.

## Results

### Design of convertible sGD elements and corresponding Cas9 lines

In previous single generation experiments, the recoded *spo11* sGD line ^12^ copied efficiently in females (76-83% transmission) and to a lesser degree in males (65-70% transmission), and drove efficiently to a high level of stable introduction in multi-generational cage experiments (~85%). Because this drive performed well but not perfectly, we selected it to test for conversion to an fGD by hacking (Figure 1a) since it would permit comparison of sGD versus fGD performance parameters in single generation crosses and in multi-generational cage trials. As a first step, we generated Cas9 donor lines, in which *vasa* or *nanos* (*nos*)-driven Cas9 transgenes are placed between a partial non-functional fragment of a tdTomato (tdTom, expressed in the eye) marker and a fully active EGFP transgene (expressed in the abdomen) that act as homology arms (HA) adjacent to a gRNA cleavage site in the sGD element (Figure 1b, *spo11* line). As detailed in Figure 1b, the sGD carries a synthetically-designed PAM site (fauxPAM) between the two marker sequences, one of which is complete and active for the sGD line (tdTom) whereas the other one is partial and inactive (EGFP) and provides the recognition site for the hacking gRNA carried outside of the homology arms on the Cas9-donor element (Figure 1b, Hacked line). The donor Cas9 transgenes, endowed with the potential to hack sGDs and referred hereon as Cas9Hack, were inserted into available AttP sites located in either the II or III chromosomes via phiC31 recombination ^18^ (Figure 1c). In addition to the aforementioned Cas9, HA, and attB sequences, Cas9Hack contains a gRNA (gRNAHack) that targets the fauxPAM site in the sGD line and mediates Cas9 insertion into the locus. gRNAHack sequences are located outside the HA-flanked Cas9 transgene, however, and thus are not copied into the sGD recipient construct (Figure 1b). When the tdTom-marked sGD and EGFP-labeled donor Cas9 elements are combined, gRNAHack-induced cleavage of the sGD element can occur. Successful conversion leads to the insertion of the Cas9 transgene plus the 5’ portion of the EGFP transgene directly upstream of the 3’ portion of the EGFP transgene into the sGD, conferring a dual fluorescence phenotype on the “hacked” element (3XP3-tdTom + Opie2-EGFP; Figure 1b, Hacked line) and bestowing autonomous self-mobilizing capacity on the composite element.

**Figure 1.**
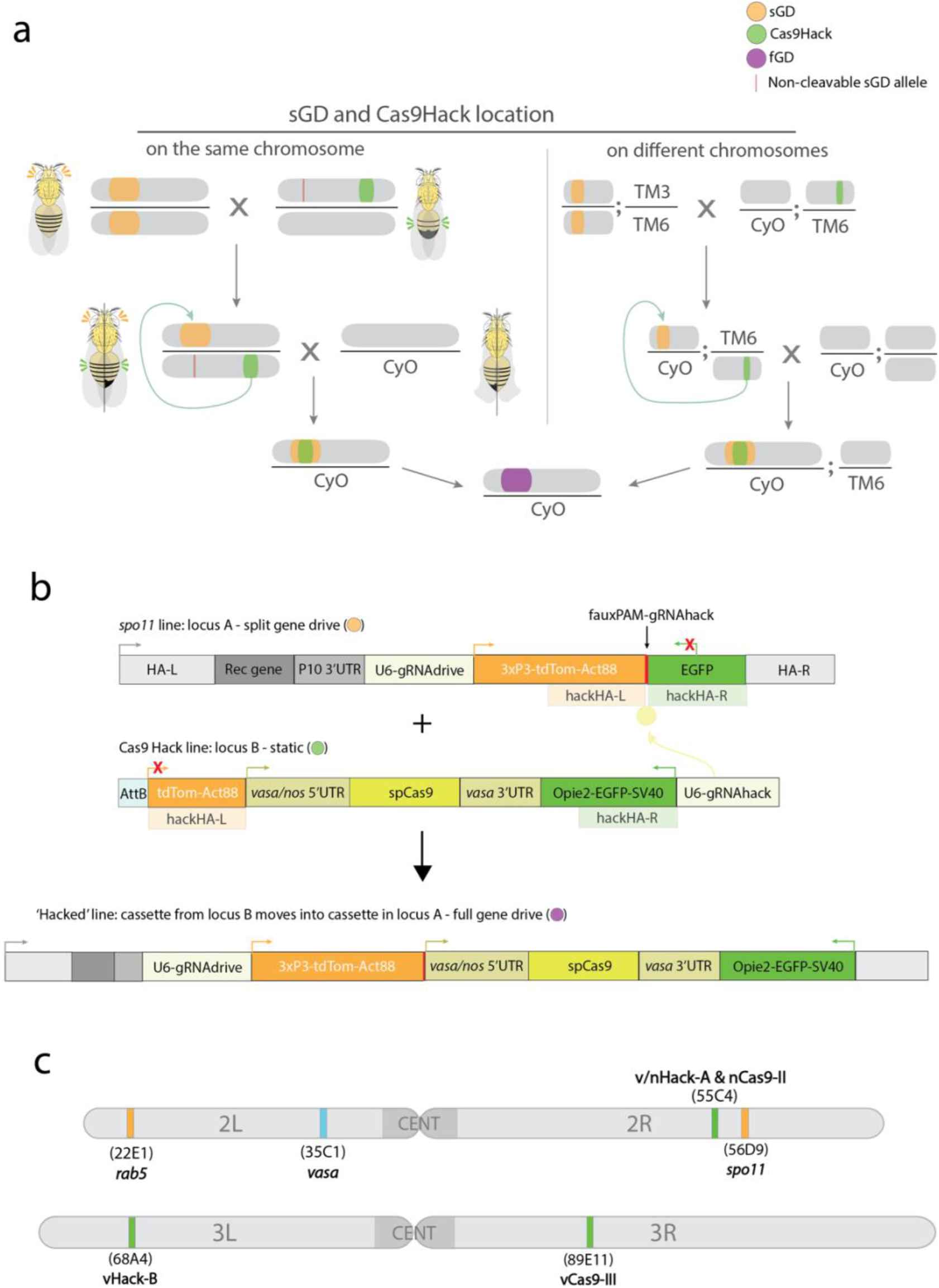
Experimental design of the split to full gene drive conversion. **a)** Outline of the genetic cross schemes used to convert a split system into a full gene drive **b)** Schematic of the genetic constructs engineered and tested in the study for sGD to fGD conversion. A tdTom-expressing split gene drive cassette (sGD) that contains a fauxPAM in between its markers is genetically paired with a specific EGFP-expressing Cas9, which contains a gRNA (that does not home) targeting the sequence next to the fauxPAM in sGD, that drives itself into the sGD locus and forms an autonomous full gene-drive cassette (fGD) using marker sequences as homology arms. The resulting cassette expresses both tdTom and EGFP markers **c)** Chromosomal location of the sGDs tested for conversion and hacking-Cas9 transgenes in the *Drosophila melanogaster* genome.

### Cas9Hack and control Cas9 lines sustain comparable copying of a sGD inserted at the *vasa* locus

In our prior study ^12^, we employed two Cas9 sources (vCas9-III and nCas9-II) expressed under the control of different promoters (*vasa* and *nos*, respectively) to drive copying of the *spo11* sGD. These two autosomal static Cas9 sources, inserted into chromosomes III and II, respectively, served as reference controls for assessing the activity of Cas9Hack sources controlled by the same promoters (vCas9Hack and nCas9Hack). The Cas9Hack transgenes were inserted in two genomic locations, 55C4 on chromosome arm 2R (Cas9Hack-A) and 64A4 on 3L (Cas9Hack-B).

In initial comparative experiments, we tested the newly generated Cas9Hack-A lines for their ability to support super-Mendelian inheritance of a reference unhackable mCerulean-marked split drive element inserted in the *vasa* locus on chromosome II (vCC; Figure 1c, Figure 2a). We chose not to use the fauxPAM-bearing *spo11* sGD to test Cas9Hack lines to avoid complications arising from gRNAHack cutting the target element simultaneously with allelic conversion and copying of that element (via its own gRNA). Homozygous G0 virgin vCC females were mated to Cas9Hack males to obtain flies trans-heterozygous for both elements (Figure 2a). Single F_1_ heterozygote males or virgin females were then crossed to wildtype flies (WT) of the opposite gender and their F_2_ progeny were scored for presence of the blue and green fluorescent markers carried by the vCC and Cas9Hack elements, respectively (Figure 2a). We observed comparable allelic conversion efficiencies for vCC using either the traditional *vasa* and *nos*-driven Cas9 (vCas9-III and nCas9-II) sources and Cas9Hack transgenes when expressed using the corresponding promoters (vHack-A and nHack-A; Figure 2a). When allelic conversion occurred in females, both traditional Cas9 lines performed comparably (vCas9-III=78±11%; nCas9-II=77±5%). Similar transmission values were also observed for Cas9Hack-A lines (vHack-A=78±9%; nHack-A =82±10%), which did not differ significantly from the control rates of vCas9-III and nCas9-II-mediated copying (two-tailed Mann-Whitney test for *vasa*: *U*=494.5, p=0.99; *nos*: *U*=170.5, p=0.07). F_2_ progeny derived from heterozygous vCC/Cas9 males displayed somewhat lower conversion frequencies as previously observed for several split drive systems in *Drosophila*, which may be related to the anomalous absence of male recombination in this species ^11^. Again, however, no significant differences were observed between Cas9 sources (vCas9-III=68±8%, nCas9-II=66±7%, vHack-A=67±10%, nHack-A=68±10%) (two-tailed Mann-Whitney test for *vasa: U*=233, p=0.91; *nos*: *U*=195, p=0.94). In all cases, due to the static nature of the tested Cas9 sources, inheritance of the Cas9 transgenes approximated the expected Mendelian frequencies (average among all conditions: 51±8%; Figure 2a).

**Figure 2.**
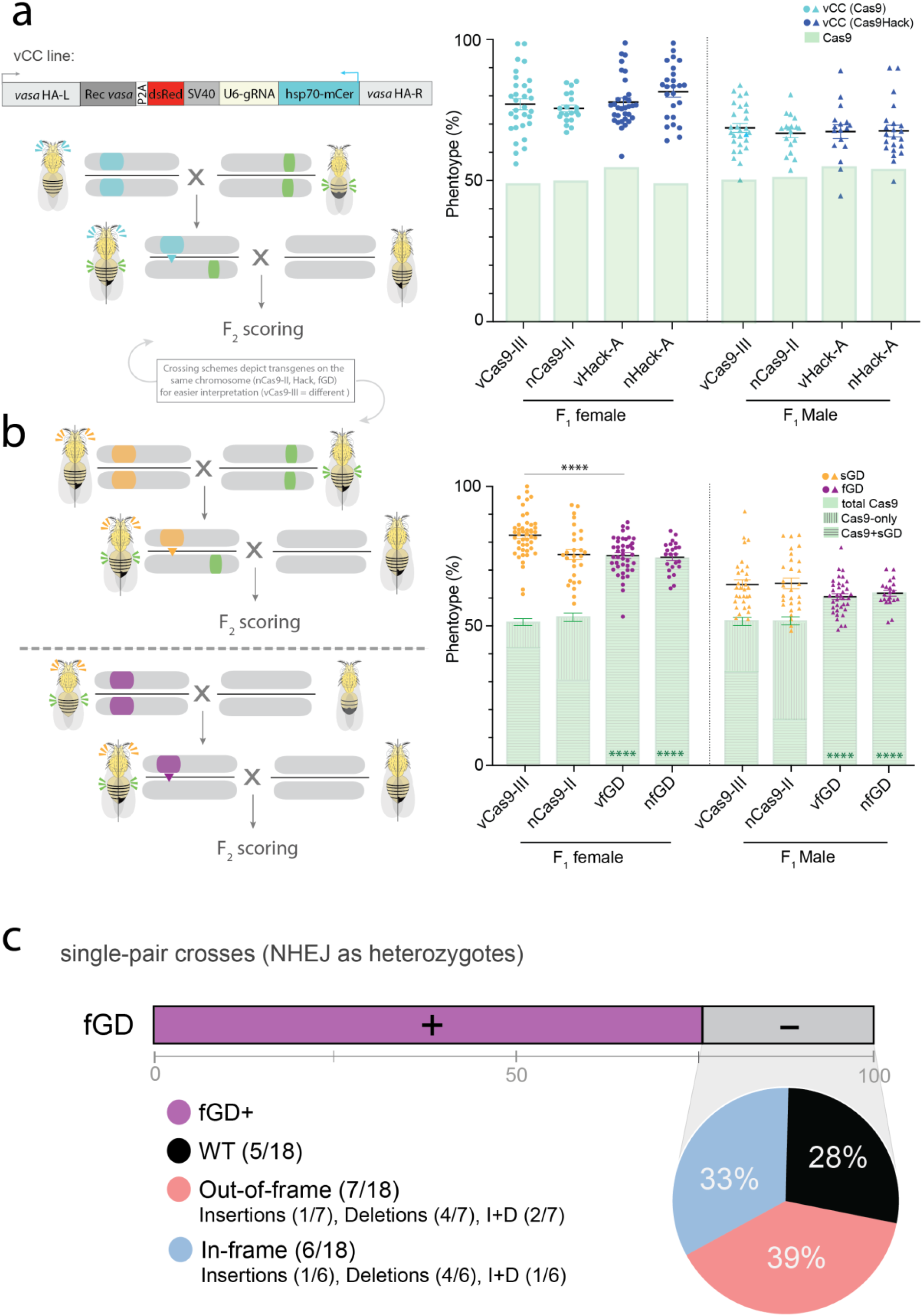
Super-Mendelian performance is comparable between split and full gene-drive elements in single-generation crosses. **a)** Performance of hacking *vasa* and *nos*-driven Cas9 was tested and compared to previously used static Cas9 lines (all marked with EGFP) by crossing them to an unhackable split drive element (vCC) inserted in the endogenous *vasa* locus, marked with mCerulean. Single F_1_ germline conversion was assessed by scoring the markers for both transgenes in the F_2_ progeny. Independent inheritance of Cas9 and vCC is depicted using green bars and blue dots, respectively. **b)** Upon successful hacking of the *spo11* sGD, *vasa* and *nos*-driven fGDs (vfGD and nfGD, respectively) were assessed for F_1_ germline conversion capacities. The panel follows the layout described in a), except for the sGD marker being tdTomato and represented in orange and purple for sGD and fGD configurations, respectively. **a, b)** Sex of the parental (F_1_) trans-heterozygote is indicated in the X-axis under the Cas9 line used, as well as by circles (female) or triangles (male), used to show the data of each individual cross. Error bars represent mean values ± SEM. Stars represent statistical significance (\*\*\*\**p* < 0.0001) on sGD-copying differences (black, two-sided *t*-test) and Cas9 deriving from Mendelian frequencies (green, *χ*^2^). Raw phenotypical data is provided as Supplementary Data. **c)** Regions surrounding the *spo11* gRNA target site were amplified from single non-fluorescent F_2_ individuals generated in b), sequenced by Sanger and analyzed. A bar depicts the % of fGD^+^ (purple) and % of non-fluorescent (fGD^−^, gray) flies. Genotypes obtained are depicted in a pie chart showing prevalence of indel mutations for the *spo11* locus. Each section of the pie chart describes the kind of NHEJ allele that is formed and % among the total tested fGD^−^ (NHEJ/WT) heterozygotes.

### Generation of *spo11* fGD lines

The comparable performance of Cas9Hack and reference Cas9 lines suggested that these transgene donor constructs were suitable for converting sGDs into fGDs by hacking. We crossed the hackable *spo11* sGD with both vHack-A and nHack-A to obtain *spo11* fGDs (Figure 1a). Of the 6 crosses we set up for each hacking element, we recovered double-marked/CyO individuals in 75% of them (5/6 for vHack-A, 4/6 for nHack-A). The predicted genomic structures of *spo11* hacked cassettes were verified for fGD elements (hacked with *vasa*: vfGD or *nos*: nfGD) using PCR amplification followed by Sanger sequencing. In contrast to recovery of these multiple independent hacking events, an identical vHack construct inserted at a different III-chromosome AttP site (vHack-B) did not produce any successful recombination events, underscoring the previously described role of genomic position effects on gene conversion efficiencies ^16^ (Figure 1c). Since the *spo11* sGD is closely linked to the vHack-A insertion site while the vHack-B locus is unlinked, the more efficient performance of vHack-A relative to vHack-*B* is consistent with prior observations of more efficient hacking from proximal loci and Hi-C sequence analysis in which neighboring segments of the genome tend to lie closer together ^19^. Consistent with this proximity hypothesis, the vHack-A (2R), which efficiently sustained hacking of the closely linked *spo11* sGD did not do so when coupled with the more distant *rab5* sGD (2L).

### *spo11* sGD allelic conversion rates are slightly higher than for the unitary *spo11* fGD element

Having successfully obtained several hacked *spo11* fGDs carrying either the *vasa*-Cas9 or *nos-* Cas9 transgenes, we next compared their allelic conversion rates to those obtained with the bipartite *spo11* sGD system. We crossed G0 homozygote *spo11* fGD males to homozygous WT virgin females and scored the fraction of fluorescently-labeled offspring (Figure 2b). In both sets of crosses, Cas9 was carried through males to avoid potentially confounding maternal transmission of the endonuclease through the female germline, although we note that attenuated carryover of this type could occur systemically in cells of fGD males due to inheritance from their grandmothers ^6,20^. Emerging F_1_ heterozygote flies were separated by gender, crossed with homozygous WT individuals of the opposite sex, and corresponding F_2_ progeny were scored for presence of the red (tdTomato) and green (EGFP) eye fluorescent markers associated with the *spo11* and Cas9 transgenes, respectively (Figure 2b). Paralleling our observations with the *spo11* sGD, where transmission through females (vCas9-III sGD=83±8%; nCas9-II sGD=76±10%) was greater than through males (vCas9-III sGD=65±9%; nCas9-II sGD=65±10%; Figure 2b), we detected gender-specific differences in both vfGD and nfGD. Overall transmission rates of fGDs from F_1_ to F_2_ progeny were approximately 75% (vfGD=75±7%, nfGD=74±6%; Figure 2b), slightly lower than those detected for the split sGD form in the case of the vCas9 source, but with comparable efficiencies observed between the fGD and sGD systems for nCas9. As mentioned above, transgene inheritance was substantially lower when passed through males, with only ~60% of the progeny (vfGD=60±6%, nfGD=62±4%; Figure 2b) receiving the transgene element, a decrease of 5% relative to both the vCas9-III and nCas9-II sGD counterparts. These modest differences between sGD and fGD transmission efficiencies in both genders may reflect different sizes of the elements given that the fGD carries an additional 7kb of cargo comprising Cas9 and its regulatory sequences. Low levels of residual Cas9/gRNA ribonucleoprotein complexes inherited from fGD grandmothers might also contribute to this slight attenuation. Alternatively, the chromosomal location of Cas9 transgenes may be relevant since Cas9 insertions on the third chromosome seemed to provide more efficient copying than those on the second chromosome. As expected, the static Cas9 sources were inherited at Mendelian frequencies in all of the *spo11* sGD crosses (vCas9-III=52±8% (both genders); nCas9-II=53±9%(F), 52±6%(M)) whereas Cas9 remained linked to the *spo11* element in the fGD (see super-Mendelian inheritance frequencies above; Figure 2b).

As a complementary approach to assess fGD performance, we examined the *in vivo* cleavage efficiency of the *spo11* fGD transgene by profiling the range of different NHEJ alleles generated in single-pair crosses (Figure 2c). We extracted DNA from non-fluorescent F_2_ offspring (fGD^−^) derived from independent crosses of an F_1_ trans-heterozygote female with a WT male. Following amplification and sequencing of the target sites in single fGD^−^ F_2_ individuals, we estimated the frequencies of non-cleaved or NHEJ-induced alleles that were transmitted to the F_2_ progeny from F_1_ trans-heterozygous mothers by correcting for the father’s WT allele from the Sanger sequencing reads. Analysis of 18 different fGD^−^ flies revealed that 28% of the individuals inherited WT sequences, while 33% carried *in-frame* indels, and 39% had *out-of-frame* mutations (Figure 2c). Based on the sequencing data and super-Mendelian conversion frequencies observed in single crosses, we estimate the overall cleavage frequency for the *spo11* fGD to be ~93%. This is very similar to the 92.3% cleavage frequency obtained previously for *spo11* as a sGD ^12^, confirming that cleavage efficiencies are locus-specific and not dependent on the size or nature of the transgenic cargo inserted at that locus.

### *spo11* sGD and fGD differ in initial cage drive dynamics but not in final outcome

In prior studies, we paired the *spo11* sGD with the vCas9-III source in cage trials seeded with initial ratios of 25% heterozygous (sGD/+; vCas9/+) to 75% WT (+/+; +/+) individuals ^12^. In those experiments, the frequency of the sGD increased rapidly to 70-85% introgression after 5-6 generations and then remained at a stable plateau due to the accumulation of non-deleterious (i.e., *in-frame* and presumably functional) NHEJ alleles (Figure 3a, orange traces). In these and other sGD experiments, we also observed that the static (or Mendelian) sources of Cas9 were eliminated from the population, which in the case of the *spo11* sGD occurred after 15 generations (Figure 3a, green traces). Since Cas9 sources on their own or in combination with other sGDs did not disappear with similar kinetics, we inferred that their loss with certain sGDs (*spo11* included) was due to fitness costs associated with carrying both sGD and Cas9 transgenes under conditions where uncleaved target alleles remained abundant.

**Figure 3.**
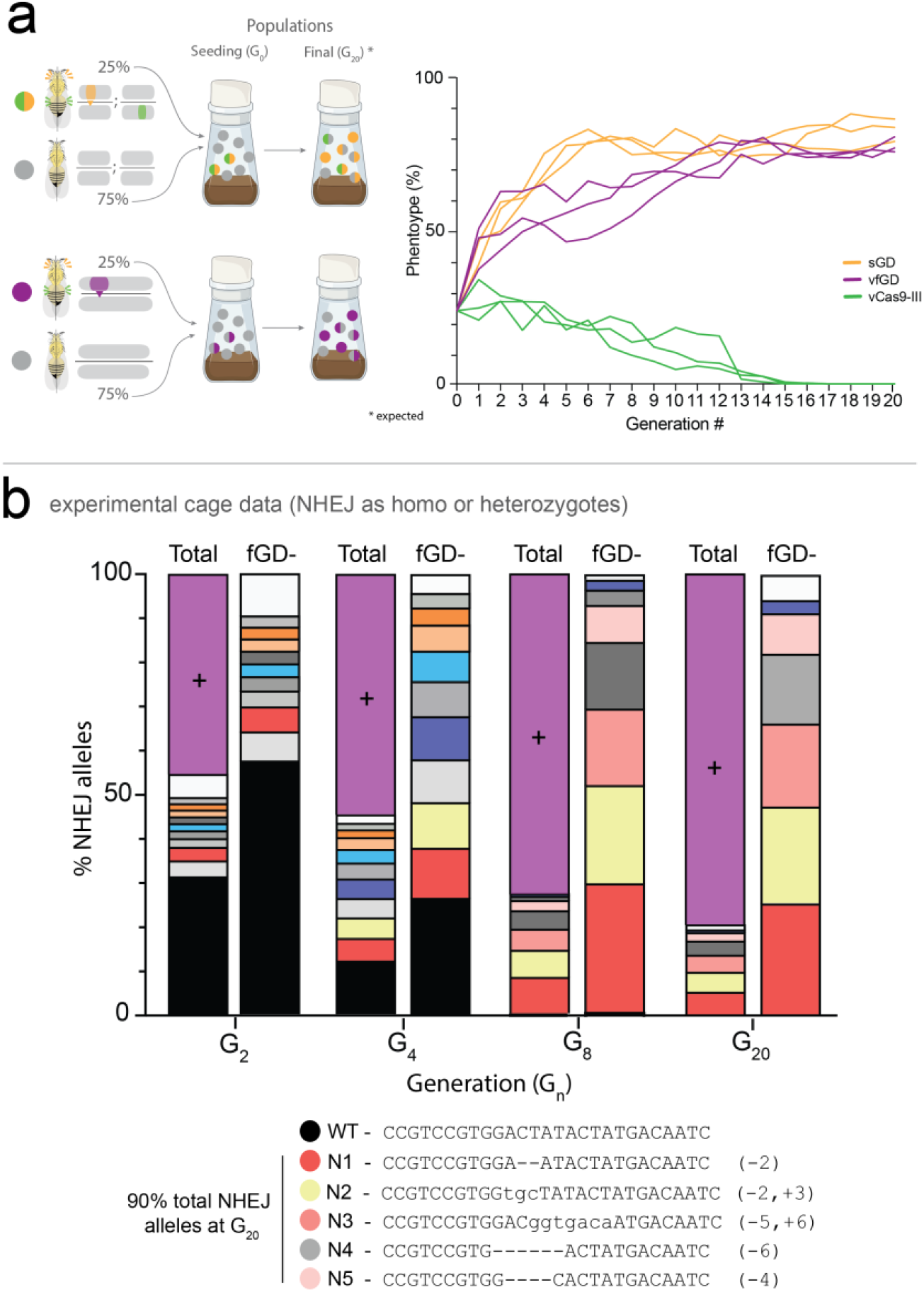
*spo11* fGD multi-generational cage trials and NHEJ profile assessment. **a)** Setup of fGD cage trials followed previous experiments with the sGD-Cas9 configuration. Virgin heterozygote fGD/+ and WT (+/+) flies were seeded at 1:3 ratio in the initial generation and allowed to mate at random at each generation (Gn). Flies in a cage were counted and scored for presence or absence of the phenotypic markers and randomly passed onto the following generation (Gn+1). Orange and green traces depict sGD cage experiments where transgene and Cas9 are unlinked and sort independently of one another. Purple traces depict fGD cage trials, where transgene and Cas9 are linked as one genomic unit. **b)** NHEJ cage trial data was obtained by deep-sequencing the target site region of pooled non-fluorescent individuals at specific generations. Most prominent NHEJ alleles at each generation are shown in bars to represent their distribution among the total population (fGD^+^ and fGD^−^, left) or only in fGD^−^ (right). Purple bars show the fGD^+^ population percentage.

Since transmission rates of vfGD and nfGD elements were comparable and similar to those achieved by the sGDs (albeit a bit lower) in single generation crosses, we selected vfGD for further comparison to vCas9-III sGD in multi-generational cage trials, where transgenic individuals face competition for mating and resources with WT individuals. We reasoned that these comparative experiments should provide insights regarding whether forced linkage of vCas9 to the gRNA within the same fGD cassette altered introduction of the fGD into a naïve population. Paralleling the sGD cage experiments, we seeded three replicate cages with 25% heterozygous (fGD/+) to 75% WT (+/+) individuals (equally mature G0 males and G0 virgin females). Randomly-selected individuals were transferred at each generation for a total of 20 generations (Figure 3a). Following an initial rise in fGD frequency, we observed a gradual increase in the percentage of drive-bearing individuals in all cage replicates, which leveled off by approximately G_12_ (Figure 3a, purple traces). These fGD plateau frequencies remained stable over the remaining 8 generations and approximated those achieved by the sGD in prior experiments ^12^. In the interval between the G_4_ and G_10_ generations, however, the drive trajectories of the sGD and fGD diverged significantly with the sGD increasing in frequency more rapidly than the fGD.

We also evaluated the DNA sequences of non-converted fGD^−^ alleles in the population at four different timepoints: G_2_, G_4_, G_8_ and G_20_ (Figure 3b) by sampling remaining non-converted alleles at every generation, then pooling 20-25 fGD^−^ individuals and performing deep-sequencing on the PCR-amplified regions flanking the Cas9 cut site. Sequence data revealed an array of different mutations at G_2_ and G_4_, but also large proportions of untouched WT alleles (Figure 3b, black bars). As was also the case of the sGD cage experiments ^12^, we observed the progressive simplification of an initially complex array of mutations generated early during the drive process, most likely due to associated fitness costs. By G_8_, WT alleles were no longer detected, presumably because they had already been acted on by Cas9 carried by the fGD^+^ allele and either successfully converted to carrying the fGD^+^ element or mutated to cleavage-resistant and largely functional NHEJ alleles. As mentioned above, the variety of NHEJ alleles decreased over time leveling out by G_8_ and remaining stable through G_20_ (except for an allele represented in grey shading for which an extra 1bp deletion appeared between G_8_ and that of G_20_). Consistent with the disappearance of convertible WT alleles by G_8_ there was only a minimal increase in the frequency of the fGD^+^ at G_20_.

### Mathematical models capture the major drive dynamics trends in population cages

In our previous studies we modeled key drive parameters, including fertility fitness costs, for the *spo11* sGD drive ^12^. Building on those models, we extend our analysis to the fGD by assessing two potential fitness cost implementations: 1) an active cost associated with ascending phase of the drive trajectory, where fitness costs due to Cas9/gRNA were only apparent during active cleavage events, and 2) a co-occurrence cost, where the fitness costs manifest whenever an individual has Cas9 and gRNA, regardless of active cleavage. Drive performance was estimated from respective cage trial data using a Naive-Bayes Multi-Objective Hidden Markov Model optimized with an evolutionary algorithm (Figure 4a). Initial parameter estimates for transmission were taken from single-pair mating data (Figure 2b) and fitness costs from the previous study ^12^, but with wider ranges to account for possible changes in fitness (see Mathematical Supplementary for a complete description of inheritance implementation, HMM, likelihood function, and estimated parameters). Cleavage rates approached 95% (95% quantiles: 90-95%, see Mathematical Supplementary Tables S11 and S12), with significantly higher HDR in females (49%, 95% quantiles: 40-58%) than males (20%, 95% quantiles 20-37%). Cleavage and conversion rates correspond to transmission estimates of 73% in females (95% quantiles: 68-78%) and 60% in males (95% quantiles: 59-68%), consistent with single-pair mating observations (Figure 2b).

**Figure 4.**
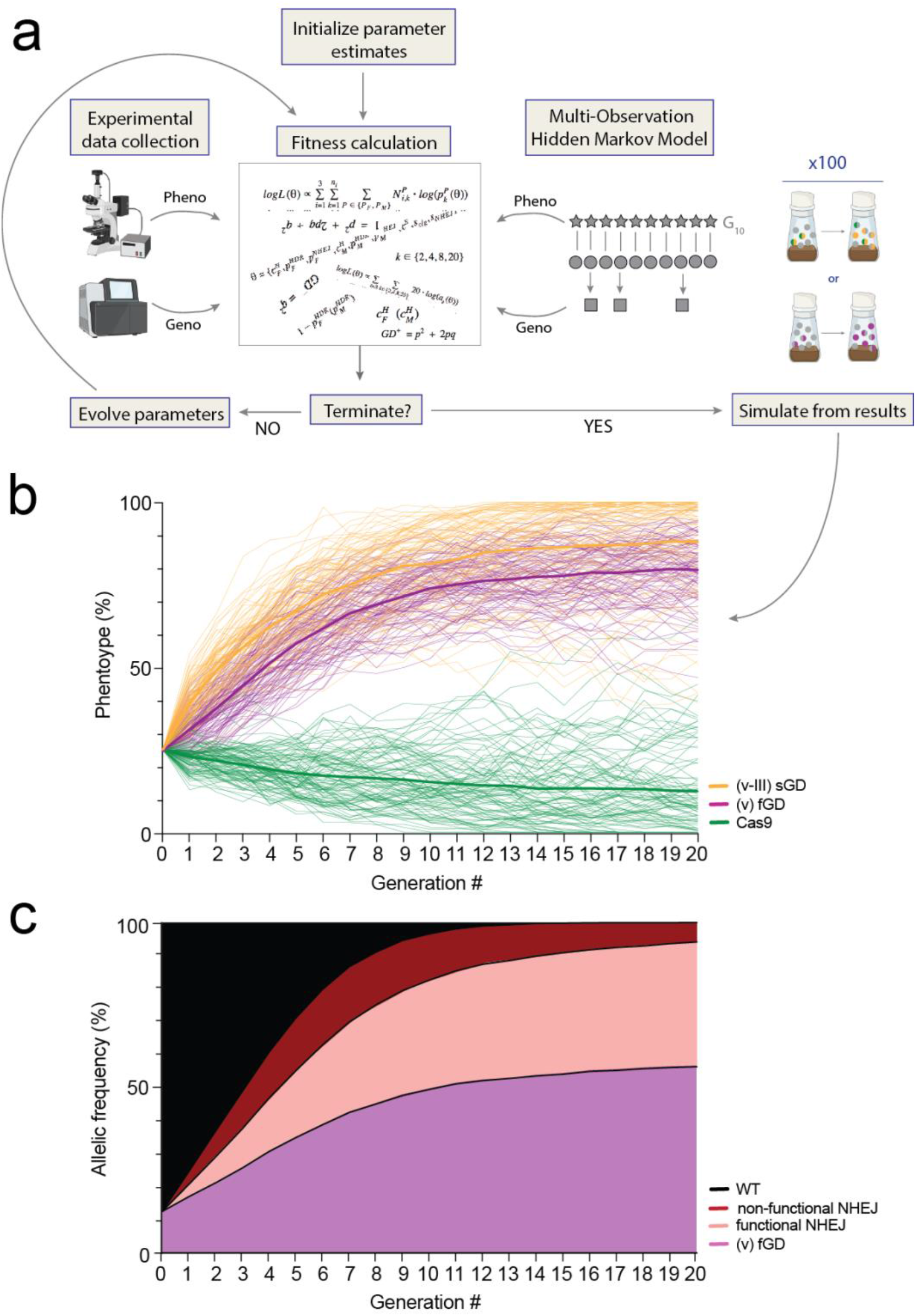
Mathematical model simulations provide insights in dynamics. **a)** Schematic of the workflow to obtain accurate mathematical models for *spo11’s* sGD and fGD configurations and their associated fitness parameters. **b)** Models were run using fitted parameter values and 100 stochastic simulations plotted for *spo11* fGD (purple) and compared to *spo11* sGD (orange for gRNA; green for Cas9). Thicker lines depict the mean of the 100 simulations. **c)** Predictions of alleles present in the fGD cage simulations at each generation. Purple, red shades and black curves depict fGD^+^, different types of NHEJ and WT allele, respectively.

Rates of *in-frame* vs *out-of-frame* NHEJ events varied widely, but preferentially towards higher *in-frame* rates, consistent with our previous work and the importance of *spo11* for fertility. However, the models lean towards no fitness cost from Cas9 and gRNAs, under either the active or co-occurrence models, which does not correlate with what was observed for the *spo11* transgene acting as sGD. This limitation of the modeling can be explained by virtue of the hacked design; the initially separated fluorescent markers from the sGD and Cas9 constructs are now linked in a single cassette (fGD), effectively presenting a single fluorescent marker than can only distinguish GD^+^ versus GD^−^ phenotypes. Thus, we applied the estimated fitness costs of Cas9 and gRNAs for the *spo11* transgene as sGD and obtained similar parameter estimates. Additionally, we used sequence data mentioned above from a sample GD^−^ flies (Figure 3c) to estimate the GD^−^ allele frequencies (see SI File, Mathematical, section Data Preparation). When considering the modest amount of sequencing data relative to the phenotypic assessments (~80 flies sequenced, compared to 19,386 flies counted for phenotype data), our models are heuristically equivalent to the measured allele frequencies with differences only minimally impacting our parameter estimates.

We generated stochastic realizations of our model, employing the active Cas9/gRNA cost and parameter estimates (see SI File, Mathematical, Tables S8 and S10, and Figure S1 for sGD and Tables S9 and S12, and Figure S3 for fGD), for comparison with the experimental cage data (Figure 4b and c). The stochastic model captured the potential role of chance events such as mate choice (multinomial-distributed), egg production (Poisson), progeny genotype (multinomial), and the finite sampling of the next generation (multivariate hypergeometric). Stochastic model trajectories were consistent with the cage trials but slightly below the long-term behavior of the sGDs (Figure 4b), likely due to the build-up of NHEJ alleles during the early generations in fGD cages corresponding to the drive delay. Additionally, allele frequencies (Figure 4c) indicate continued clearance of *out-of-frame* NHEJ alleles, consistent with winnowing deleterious alleles of an essential gene and with sequencing results obtained (Figure 3b).

## Discussion

In this study, we genetically converted a split gene-drive element targeting a locus essential for fertility in both sexes (*spo11*) into a full gene-drive. The sGD element was designed to contain homology arms flanking a Cas9 transgene that acts as a donor template for insertion into the sGD upon cleavage of the locus. We observed that sGD-to-fGD conversion efficiency depends greatly on the proximity between Cas9-Hack elements and target sGD loci. We tested two Cas9 transgenes (vHack-A or nHack-A driven by *vasa* or *nos*, respectively) that were closely linked (7-10 cM) to the sGD (2R) and assessed their capacity to copy into a suitable transgene located in the *spo11* locus. Both *vasa* and *nos* Cas9Hack-A transgenes supported sGD-to-fGD conversion at high frequencies, as we observed multiple hacking events (3 of every 4 crosses). However, when testing the exact same hacking Cas9 donor constructs (CasHack-B) located on a different chromosome (3L) or an alternative freely recombining target sGD line (*rab5*) located in the opposite chromosome arm (2L) in combination with the vHack-A line homologous recombination events were not recovered. These observations agree with those described previously ^17,21,22^, as chromosomal repair or HACK efficiency varied depending on genomic distance and orientation of the target-donor pair. Hi-C distances are highly relevant in this context, as they estimate the actual physical DNA proximity between two loci within meiotic nuclei. Thus, in future efforts to convert additional sGDs to fGDs, Cas9Hack lines inserted near the genomic integration site of the sGD element in question may prove most efficient.

Following successful recovery of both *spo11* vfGD and nfGD lines, we assessed their copying performance in single generation crosses and compared them to that of their non-autonomous sGD counterparts. Paralleling outcomes observed with sGD crosses, all fGD-carrying females displayed a greater average conversion efficiency (75%) than males (62%) as has been generally observed for autosomal drives in *Drosophila* ^11,12,23^, which may be related to the anomalous absence of male recombination in this species. In single generation crosses, the *spo11* fGD lines copied at slightly lower rates (~5-7%) than the sGD. The chromosomal position of the Cas9 transgene relative to *spo11*, or the greater overall size of the fGD element, may contribute to these modest reductions in conversion efficiencies. Regarding the potential role of genome position effects for Cas9 transgene insertion, we compared all Cas9 lines against a non-hackable sGD, *v*CC, and observed that the inheritance of the transgene was very similar for all of them. While chromosomal position of the tested Cas9 sources did not matter in this instance, Cas9-dependent drive performance has been noted in other sGD-Cas9 combinations ^12,14^. Cassette size may also influence conversion efficiencies, which could also be relevant in the context of the fGD element as the hacking mechanism inserts the whole promoter-Cas9 cassette into the locus, increasing the fGD cassette size to ~8kb longer than the sGD. Reduced copying efficiency has also been observed for larger drives inserted into the same *yellow* locus ^11,24^.

We also tested the performance of the *spo11* vfGD in multi-generational cage trials where we observed that it had the capacity to spread through a naïve population, as expected, following initial introduction at low levels. When seeded at a frequency of 12.5% of the total alleles (phenotypic: 25%), the vfGD phenotype increased steadily to ~80% of the population in 15 generations. This steady-state level corresponds to a similar degree of final introgression, but with delayed drive kinetics, compared to cage studies with the *spo11* sGD paired with the corresponding *vasa*-Cas9 source. In that case, we observed a similar increase in total frequency, but in approximately half the number of generations ^12^. Final levels of introgression were comparable for the two systems since they both plateau after reaching a phenotypic frequency of 75-85%, mostly due to the generation of a similar number and complexity of non-deleterious resistant NHEJ alleles. In the previous study ^12^, we observed that there was a moderate fitness cost associated with carrying certain sGDs and a Cas9 transgene in the same individual. In contrast to the sGD configuration, where copying of the gRNA-bearing cassette and static Cas9 transgene can separate during transmission (due to independent assortment of the Cas9 transgene), Cas9 is forcibly linked to gRNA in the fGD and thus a higher percentage of individuals will incur costs associated with co-inheritance of these CRISPR components. This effect is particularly pronounced in early generations when the great majority of non-transgenic individuals will be homozygous for the wildtype genotype and thus have fitness advantage over the transgenic line during the drive process. Since fitness costs associated with co-inheritance of gRNA and Cas9 transgenes are confined to the active drive phase (due to a strong form of lethal or sterile mosaicism acting on wild-type paternal alleles), the fGD eventually achieves levels of introgression comparable to the sGD and then remains at stable levels in the population as nearly all remaining alleles are cleavage resistant (and most likely functional).

Mathematical modeling confirmed that many of the inferred drive parameters are similar for the sGD and fGD population cage experiments. Such modeling is rapidly becoming requisite in the gene drive toolbox, both for the analysis of experimental data ^24,25^ and simulation of potential results to inform conservation ^26^ and disease-elimination campaigns ^27,28^. In this work, we demonstrate the application of a Naive-Bayes multi-observation HMM that optimizes the regular HMM in a highly-parallel framework by incorporating phenotypic and genotypic data. Simulations using our models (Figure 4) recapitulate what we observed in experimental data and provide us with ranges of plausible fitness cost values and parameters (Supplementary Information file, Mathematical).

In summary, this study provides a direct comparison between split and full drive configurations and validation of a general method for efficiently converting the one to the other by genetic means. These studies reveal unexpected differences in the kinetics between the split versus coupled drives in which the former displayed more rapid initial drive than the full drive (most likely due to a transient fitness cost of obligate co-inheritance of Cas9 and gRNA transgenes) but then reach comparable levels of final introgression. Thus, both systems offer potential advantages for field applications. The split system provides the potential for more localized and controllable spread of the CRISPR components while full gene drives could be implemented in contexts where long-lasting protection conferred by associated effector genes may be desired.

## Materials and Methods

### Plasmid construction

All plasmids were cloned using standard recombinant DNA techniques. Plasmid and genomic DNA sequences were amplified using Q5 Hotstart Master Mix (New England Biolabs, Cat. #M0494S) and Gibson assembled with NEBuilder HiFi DNA Assembly Master Mix (New England Biolabs, Cat. # E2621). Resulting plasmids were transformed into NEB 5-alpha chemically-competent *E. coli* (New England Biolabs, Cat. # C2987), isolated and sequenced. Primer sequences used for the creation of the different Cas9Hack plasmids can be found in the Supplementary File. *Spo11* and *rab5* sGD plasmid construction has been described previously ^12^. Recoded cDNA fragments were designed by using non sub-optimal alternative codons from CRISPR cut site to gene’s stop codon and synthesized as gBlocks™ (Integrated DNA Technologies). Codon usage was kept as similar as possible to that of the endogenous sequence.

### Microinjection of constructs

Plasmids were purified using the PureLink Fast Low-endotoxin Maxi Plasmid Purification kit (ThermoFisher Scientific, Cat. #A35895). All plasmids were sequenced prior to injection. Embryo injections were carried out at Rainbow Transgenic Flies, Inc. (http://www.rainbowgene.com). Each Cas9Hack construct was injected into AttP-harboring lines expressing integrase in the X chromosome (Bloomington #R8621 and #R8622). Injected embryos were received as G0 larvae, allowed to emerge and 3–4 females were intercrossed to 3–4 males. G_1_ progeny were screened for positive transgene marker (green body). All transgenic flies that displayed the marker were then balanced using Sco/CyO (for Cas9Hack-A, located on the II chromosome) or TM3/TM6 (Cas9Hack-B, III chromosome) and kept on a w^1118^ background. Homozygous stocks were kept in absence of any balancer alleles or markers associated to the initial inserted line. Correct and complete transgene insertions in homozygous stocks were validated through PCR amplification and Sanger sequencing.

### Fly genetics and crosses

Fly stocks were kept and reared on regular cornmeal medium under standard conditions at 20–22 °C with a 12-hour day–night cycle. sGD and Cas9Hack stocks were kept separate in glass vials in an ACL-1 fly room, freezing the flies for 48 h prior to their discard. To assess Cas9Hack lines ability to drive, we genetically crossed vCC to each Cas9Hack line. To do so, individual trans-heterozygote F_1_ males or virgin females were collected for each G0 cross and crossed to a wildtype fly of the opposite gender. Single generation crosses were grown at 25 °C. Inheritance of both gRNA (>50%) and Cas9 (~50%) were calculated using the resulting F_2_ progeny by scoring the phenotypic markers associated to each transgenic cassette. Hacking efficiency experiments and all experiments performed after successful Cas9 introduction into the *spo11* sGD locus, *spo11* Hack lines were maintained in plastic vials in a contained ACL-2 insectary dedicated to *Drosophila* gene drive research. Used vials or samples were frozen for at least 48h prior to their removal from the facility to be discarded or worked with, respectively.

### Multigenerational cage trials

All population cage experiments were conducted at 25 °C with a 12-hour day-night cycle using 250 ml bottles containing standard cornmeal medium. Crosses between homozygous fGD^+^ and wildtype (fGD^−^) flies were carried out to obtain F_1_ fGD^+^/fGD^−^ heterozygotes, which were used to seed the initial generation. Wildtype or heterozygote males and virgin females were collected and separately matured for 3 to 5 days. Cages were seeded at a phenotypic frequency of 25% fGD^+^ heterozygotes (15 males, 15 females) to 75% fGD^−^ (45 males, 45 females). Upon transfer into fresh bottles for each generation, flies were allowed to mate and lay eggs for 3 days, then were removed from the cage (Gn) and bottles were kept for 10 days to allow for the subsequent generation to develop to adulthood. Adult progeny (Gn+1) was randomly separated into two pools and scored; one pool collected for sequencing analyses while the other was used to seed the following generation. If the two pools differed much phenotypically, frequencies were averaged in order to reduce variability and stochastic extremes. Continuous sampling and passage was carried out for 20 generations.

### Molecular analysis of resistant alleles

To extract fly genomic DNA for single fly resistant allele sequence analysis, single flies were squashed in lysis solution (10 mM Tris-Cl pH 8.2, 1 mM EDTA, 25 mM NaCl and 0.2mg/ml proteinase K), incubated at 37 °C for 30 min and deactivated at 95 °C for 2 min. After extraction, each sample was diluted in water and stored at −20 °C if needed. 1–2 μl of each diluted DNA extraction was used as template for a 25 μl PCR reaction that covered the flanking regions of the gRNA cut site, which were used to sequence the alleles. Sanger sequencing in individual non-fluorescent flies was performed at Genewiz, Inc. in San Diego, CA to obtain the single-cross NHEJ data. NHEJ allele sequences were obtained from Sanger chromatographs by isolating the WT sequence first and then annotating the remaining allelic sequence. For cage trials, 20-25 non-fluorescent flies were pooled and their DNA extracted at each sampling generation (G_2_, G_4_, G_8_ and G_20_). Target sequences were amplified using specific gene primers that also contained adapter sequences. Non-fragmented amplicons were sequenced using Illumina-based technology (2×250bp reads, Amplicon-EZ, Genewiz), with final data being delivered as FASTQ reads and aligned to a reference sequence for each gene of interest to detect indel formation. Primer sequences used for either single fly or cage trial deep sequencing analyses can be found in the Supplementary File.

### Mathematical modeling

Model fitting was performed using a discrete-generation adaptation of the Mosquito Gene Drive Explorer (MGDrivE) ^29^ software. A multi-objective HMM, using the log-likelihood as a score function, was optimized by evolutionary algorithm and used to generate parameter estimates with corresponding 95% quantiles. Mendelian inheritance was assumed except under co-occurrence of the Cas9 and gRNA constructs, when the split-drive (and HACK) design allowed active cleavage of the target chromosome and the possibility of super-Mendelian inheritance. When cleavage occurred, a fraction of the cut alleles could be properly repaired via HDR, and the remaining cut alleles underwent NHEJ repair, generating *in* or *out-of-frame* resistant alleles. The effects of shadow-drive ^6,20^, in which Cas9 protein is deposited in the embryo of a female individual who does not carry the Cas9 allele, but whose own mother does, were also accommodated. Fitness costs were implemented as fractional reductions in male and female fecundity, consistent with *spo11* activity, and tested under active cleavage conditions as well as co-occurrence of Cas9 and gRNAs. Additional details on the model implementation and likelihood function used in sGD and fGD model fitting can be found in the Supplementary Information. Phenotype and genotype mappings are provided in Tables S2 through S7. Fitness cost implementations, active vs co-occurrence, are shown in Tables S8 and S9. Parameter descriptions, estimates, and 95% quantiles are provided in Tables S10 through S13. The resulting fits are visualized, with cage trial data, in Figures S1 through S4. Figures S5 and S6 analyze parameter estimate correlations while Figures S7 and S8 are partial rank correlation coefficient (PRCC) analyses.

Simulated model trajectories for Figure 4 were generated using a stochastic implementation of the discrete-generation model. At each generation, adult females mate with males, thereby obtaining a composite mated genotype (their own, and that of their mate) with mate choice following a multinomial distribution determined by adult male genotype frequencies, modified by mating efficacy. Egg production by mated adult females then follows a Poisson distribution, proportional to the genotype-specific lifetime fecundity of the adult female. Offspring genotype follows a multinomial distribution informed by the composite mated female genotype and the inheritance pattern of the gene drive system. Sex distribution of offspring follows a binomial distribution, assuming equal probability for each sex. Female and male adults from each generation are then sampled equally to seed the next generation, with sample size proportional to the average size of the cage trials at that generation, following a multivariate hypergeometric distribution. All simulations were performed and analyzed in R ^30^.

### Figure generation and statistical analysis

Initial graphs were generated using Prism 9 (v9.2, GraphPad Software Inc., San Diego, CA) and modified using Adobe Illustrator (v25.4.1, Adobe Inc., San Jose, CA) to visually fit the rest of the non-data figures featured in the paper. Figures 3a and 4a contain parts that were generated using BioRender.

### Safety measures

All research involving non-hackable gRNA cassettes/split gene drives was performed in glass vials in an ACL-1 facility, whereas research involving full gene drives (*spo11* Hack lines) was conducted in plastic disposable vials in an ACL-2 facility, in accordance with the Institutional Biosafety Committee-approved protocol from the University of California San Diego. All vials were frozen for 48 h prior to autoclaving and discarding the flies. ACL-2 samples were also frozen for at least 48 h before their removal of the contained facility.

## Data and code availability

Primers used for plasmid construction and sequencing (single and deep-sequencing experiments) can be found in Table S1. Full plasmid sequences can be found at the end of the manuscript’s Supplementary Information File. Raw experimental data is also provided in this paper as a Supplementary Data file. All remaining experimental data is available from the authors upon request. Modeling information can be found as a Supplementary Method of this study in the Supplementary Information file. A version of MGDrivE was used for simulation modeling and is freely available from the MGDrive GitHub repository (https://marshalllab.github.io/MGDrivE/). Specific code can be obtained from the authors upon request.

## Funding

Research was supported by a Paul G. Allen Frontiers Group Distinguished Investigators Award to E.B., NIH grant R01GM117321, and by gift from the Tata Trusts in India to TIGS-UCSD. J.B.B. and J.M.M. were supported by the DARPA Safe Genes Program Grant (HR0011-17-2-0047).

## Competing Interests

E.B. has equity interests in Agragene Inc. and Synbal Inc., companies that may potentially benefit from the research results. E.B. also serves on the company’s Board of Directors (Synbal) and Scientific Advisory Board (Synbal and Agragene). The terms of this arrangement have been reviewed and approved by the University of California, San Diego in accordance with its conflict-of-interest policies. All other authors declare no competing interests.

